# Non-antibiotic pharmaceuticals exhibit toxicity against *Escherichia coli* at environmentally relevant concentrations with no evolution of cross-resistance to antibiotics

**DOI:** 10.1101/2023.08.21.554069

**Authors:** Rebecca J Hall, Ann E Snaith, Sarah J Element, Robert A Moran, Hannah Smith, Elizabeth A Cummins, Michael J Bottery, Kaniz F Chowdhury, Dipti Sareen, Iqbal Ahmad, Jessica M A Blair, Laura J Carter, Alan McNally

## Abstract

Antimicrobial resistance can arise in the natural environment via prolonged exposure to the effluent surrounding manufacturing facilities. These facilities also produce non-antibiotic pharmaceuticals, and the effect of these on the surrounding microbial communities is less clear; whether they have inherent toxicity, or whether long-term exposure might select for cross-resistance to antibiotics. To this end, we screened four non-antibiotic pharmaceuticals (acetaminophen, ibuprofen, propranolol, met formin) and titanium dioxide for toxicity against *Escherichia coli* K-12 MG1655 and conducted a 30 day selection experiment to assess the effect of long-term exposure. All compounds reduced the maximum optical density reached by *E. coli* at a range of concentrations including one of environmental relevance, with transcriptome analysis identifying upregulated genes related to stress response and multidrug efflux in response ibuprofen treatment. The non-antibiotic pharmaceuticals did not select for significant genetic changes following a 30 day exposure, and no evidence of selection for cross-resistance to antibiotics was observed for population evolved in the presence of ibuprofen in spite of the differential gene expression after exposure to this compound. This work suggests that these non-antibiotic pharmaceuticals, at environmental concentrations, do not select for cross-resistance to antibiotics in *E. coli*.

## Introduction

Antimicrobial resistance (AMR) is a leading public health concern with global but unequal causes and consequences [1; 2; 3]. There is concern about the extent to which AMR is driven by effluent from pharmaceutical production and wastewater treatment facilities that can enter local ground and surface water [4; 5; 6; 7; 8]. This long-term supply of wastewater is known to have considerable effects on the local microbial community [8; 9; 10]. Metagenomic studies have for example identified AMR genes in water bodies close to pharmaceutical plants in countries including India and Croatia [6; 11; 12]. High levels of antibiotics and antibiotic resistance genes have also been measured around production facilities in Lahore, Pakistan [13]. Whilst the evolution of bacteria in response to antibiotics is well understood [14; 15; 16], less is known about the possible impact of long-term exposure to non-antibiotic pharmaceuticals. Importantly, manufacturing facilities typically produce more than one chemical entity, meaning waste from these sites can contain a range of biologically active chemicals including both antibiotics and non-antibiotic compounds [17].

Like antibiotics, non-antibiotic pharmaceuticals are ubiquitous contaminants that have been detected in water bodies globally following inadvertent release into the environment [5; 18; 19], and efforts have been made to begin characterising their effects on different bacterial species. Compounds including vasodilators and selective norepinephrine re-uptake inhibitors have, at various concentrations, antimicrobial activity against *Escherichia coli*, *Staphylococcus aureus*, *Pseudomonas aeruginosa*, and *Candida albicans* [20]. A screen of non-antibiotic pharmaceuticals against 40 species of gut bacteria found over 200 human-targeted drugs that had a negative effect on the growth of at least one of the species tested [21]. Interesting, the majority of these compounds were found to be active against only a few strains, suggesting that non-antibiotic pharmaceuticals may have strain-specific effects. Titanium dioxide (TiO_2_), found in a wide range of products from cosmetics to paints [22], has been shown, at various concentrations, to have antibacterial effects on organisms including *E. coli* [23; 24; 25], *P. aeruginosa* [23], *S. aureus* [23; 24; 25; 26], and *Enterococcus faecalis*, amongst others [27; 28]. In many instances, the antimicrobial properties of TiO_2_ have been evaluated as a surface coating [29; 24] rather than a suspension, the latter representing the form in which microbial communities would be exposed to TiO_2_ in waste effluent. The anti-inflammatory drug ibuprofen has been demonstrated, using disc diffusion methods, to exhibit antimicrobial activity against species including *S. aureus* and *Bacillus subtilis*, whilst not against *E. coli* or *P. aeruginosa* [30], further highlighting the species-specific activity of these compounds.

There has also been recent interest in the potential of non-antibiotic pharmaceuticals to influence the susceptibility of bacterial species to antibiotics. Compounds including ibuprofen, diclofenac, and propranolol have been linked to enhanced uptake of antibiotic resistance genes, possibly due to an increase in cell competency and membrane permeability, and the promotion of conjugative plasmid transfer [31; 32]. The antiepileptic drug carbamazepine has also been shown to promote the transfer of plasmid-encoded resistance genes via conjugation [33]. In contrast, experiments in clinically relevant species suggest that metformin, used in the treatment of type 2 diabetes, and the non-steroidal anti-inflammatory drug benzydamine can promote uptake of tetracyclines, thereby reversing a resistance phenotype in multidrug resistant pathogens [34; 35], and antibiotic-non-antibiotic drug combinations have been suggested as possible routes for treating infections caused by such species [36]. The short- and long-term effects of exposure to these compounds on bacterial populations therefore warrant further study; whether they have intrinsic toxicity and, if so, the mechanism of action and the likelihood that they could act as selection pressures resulting in significant genetic changes. Importantly, discovery of the latter might indicate a possible mechanism by which exposure to non-antibiotic pharmaceuticals could select for cross-resistance to antibiotics. This is therefore an area of research with important clinical ramifications.

Here, we undertook an investigation of the short- and long-term effects of a panel of non-antibiotic compounds on *Escherichia coli* K-12 MG1655, with an emphasis on testing at environmentally-relevant concentrations [37; 38; 39; 40; 41; 42; 43; 44] to examine the potential for selecting for cross-resistance to antibiotics. The compounds selected were TiO_2_, acetaminophen (the active ingredient in the painkiller paracetamol), ibuprofen (an anti-inflammatory), propranolol (a beta-blocker used to treat heart conditions), and metformin (a medication for type 2 diabetes). These compounds were all found to have a degree of toxicity against this strain of *E. coli* at a range of concentrations including those of environmental relevance, with transcriptome analysis identifying upregulated genes involved in stress response and multidrug efflux during ibuprofen treatment. Through experimental evolution in the presence of environmentally relevant concentrations of these compounds we found evolved populations displayed decreased fitness relative to the ancestral lineage when grown in the presence of the selection compound. However, analysis of hybrid assemblies of the evolved isolates found no single nucleotide polymorphisms (SNPs) between independently evolved populations, and there was no change in minimum inhibitory concentration (MIC) for a panel of antibiotics against isolates evolved in the presence of ibuprofen compared to the ancestor. Together, this suggests that the toxicity of the non-antibiotic pharmaceuticals does not exert a selection pressure sufficiently strong enough to lead to the fixation of mutations under the conditions tested, and with no observed selection for cross-resistance to antibiotics.

## Methods

### Strains and growth conditions

To measure potential toxicity of compounds, *E. coli* K-12 MG1655 was streaked from a glycerol stock on to a Luria Bertani (LB) agar plate (E & O Laboratories Ltd), incubated overnight at 37*^◦^*C. A single colony was then used to inoculate 5 mL LB (E & O Laboratories Ltd) in a 30 mL universal before overnight incubation at 37*^◦^*C with agitation. The overnight cultures were diluted to an optical density at 600 nm (OD600) of approximately 0.5 in LB. Serial dilutions of acetaminophen, ibuprofen, titanium dioxide (TiO_2_, in the form of nanoparticles), propranolol, and metformin were prepared as per Table S1. These compounds were selected as both published literature and preliminary investigations identified their presence in wastewater and receiving water environments, and they are commonly used non-antibiotic pharmaceuticals [5]. Additionally, TiO_2_ is found in a wide range of products and has suggested applications in water treatment [27; 45; 46]. Environmentally relevant concentrations were identified following a literature search and are provided in Table S1. A solution of 99 *µ*L of LB + compound was added to each test well of a 96-well plate, including an LB-only control, with 1 *µ*L of the dilute cell suspension then added. Plates were incubated for 24 hours in a microplate reader (Tecan) at 37*^◦^*C with continuous double orbital shaking, with absorbance measurements (OD600) taken every 30 minutes in triplicate. To assess the growth kinetics of evolved populations, ten colonies were selected for incubation as representative of the population and the kinetics monitored in a microplate reader as described previously, in the presence of the compound to which they were exposure during the selection experiment. Compounds were tested at 1x and 100x selection concentrations (Table S1) to measure whether the evolved isolates would show improved growth compared to the ancestral lineage when stressed with a higher concentration of the compound to which they had been exposed during the selection experiment.

### Genome sequencing and bioinformatics

Illumina short read sequencing of the ancestral and evolved isolates was performed by MicrobesNG (UK). Long-read sequencing of the same isolates was performed using MinION sequencing (Oxford Nanopore Technologies, UK). Briefly, genomic DNA was extracted from overnight cultures using the Monarch Genomic DNA Purification Kit (New England Biolabs). DNA was quantified using a Qubit 4 fluorometer (Invitrogen) and accompanying broad-range double stranded DNA assay kit (Invitrogen). Sequencing libraries were prepared using SQK-LSK109 ligation sequencing kit and EXP-NBD114 native barcode expansion (Oxford Nanopore Technologies, UK), as per manufacturer instructions. Long-read sequencing was performed on a MinION sequencer using an R9.4.1 flow cell (Oxford Nanopore Technologies, UK). Base calling was conducted using Guppy (v6.0.1). Reads were filtered using Filt-long (v0.2.1) using a cut-off of 600000000 target bases and demultiplexed using qcat (v1.1.0). Hybrid assemblies were then generated using Unicycler (v0.4.8-beta) in bold mode. Panaroo (v1.2.10) was used to generate gene presence/absence and core gene alignment files with the latter used to construct a maximum likelihood tree with IQ-TREE (v2.2.0.3). The tree and gene presence/absence data were visualised in Phandango [47] to look for differential gene presence patterns across the evolved isolates. A custom ABRicate (v0.8) database was used to investigate the presence and identity of the *ldrA* gene across the evolved isolates. The presence of SNPs was analysed using snippy (v4.3.6). Potential movement of insertion sequence (IS) elements was investigated using ISEScan (v1.7.2.3), ISFinder [48], and a custom ABRicate (v0.8) database, with sequences interrogated in Unipro UGENE (v47.0).

### Transcriptome sequencing

RNA sequencing was performed on *E. coli* grown in the presence and absence of 50 *µ*g/mL ibuprofen in triplicate. For the control (absence) replicates, an equivalent volume of the ibuprofen solvent (ethanol) was added. For sample preparation, a single colony for each replicate was picked following overnight growth on LB agar and added to 5 mL of LB broth (Sigma-Aldrich, UK). A 100 *µ*L suspension of each overnight culture was then transferred into 10 mL fresh LB in the presence or absence of 50 *µ*g/mL ibuprofen, with cultures then incubated at 37*^◦^*C with agitation until an optical density at 600 nm (OD600) of approximately 0.9. A 1 mL sample was centrifuged for five minutes at 10,000 rpm (Eppendorf MiniSpin F-45-12-11), resuspended in 1 mL phosphate buffered saline (PBS, VWR), and this wash step repeated. The supernatant was aspirated and the pellet frozen prior to processing and RNA sequencing by GENEWIZ from Azenta Life Sciences (Frankfurt, Germany) using their standard RNA sequencing service. Differential gene expression was quantified using Kallisto (v0.48.0) A long-read assembly of the ancestral *E. coli*, annotated using Prokka (v1.14.6), was used as a reference. The annotated assembly was processed using genbank_to_kallisto.py (https://github.com/AnnaSyme/genbank_to_kallisto.py). GNU parallel [49] was used for job parallelization. Differential gene expression was analyzed in Degust (v4.1.1) with a false discovery rate threshold of p *<* 0.05 and an absolute log fold change of at least 1.

### Selection experiment

The ancestral *E. coli* isolate was streaked from a glycerol stock on to an LB plate and incubated overnight at 37*^◦^*C. A single colony was used to inoculate 5 mL nutrient broth (NB) (Sigma) in a 30 mL universal, with six independent biological replicates per condition. Acetaminophen (5 ng/mL), ibuprofen (2 *µ*g/mL), TiO_2_ (1 *µ*g/mL), propranolol (0.5 ng/mL), and metformin (0.5 ng/mL) were tested individually, including a NB-only control. Microcosms were incubated for 24 hours at 37*^◦^*C with agitation, before a 1% transfer of cell suspension into fresh media. This 1% transfer was repeated every 24 hours for 30 days. After 30 days, the whole population was centrifuged at 3600 rpm (Thermo Scientific Megafuge 40R TX-1000) for five minutes, resuspended in 1 mL 50% glycerol, and stored at −80*^◦^*C. To assess whether the populations were experiencing short-term, reversible toxicity as a result of compound exposure, ten colonies from each end-point population were selected from a UTI ChromoSelect agar plate (Millipore) and used to inoculate 5 mL NB only. The microcosms were incubated for 24 hours at 37*^◦^*C with agitation, before a 1% transfer of cell suspension into fresh media every 24 hours for seven days. After seven days, the whole population was centrifuged at 3600 rpm (Thermo Scientific Megafuge 40R TX-1000) for five minutes, resuspended in 1 mL 50% glycerol, and stored at −80*^◦^*C.

### Minimum inhibitory concentration assay

Stocks of ethidium bromide, ampicillin, ciprofloxacin, chloramphenicol, trimethoprim, and colistin were prepared to 1000 *µ*g/mL. Cultures of the *E. coli* MG1655 ancestor and an *E. coli* ATCC 25922 control strain were prepared by inoculating 5 mL of LB with a single colony of bacteria and incubating with agitation at 37*^◦^*for 18 hours. Iso-Sensitest broth (ISB) (Thermo Fisher Scientific) was then used throughout the assay. The overnight culture was then diluted 1:100 and working stocks of antibiotics prepared, both in ISB. U-bottom 96-well plates were set up so that the cultures were incubated with no antibiotic and with 11 different concentrations of antibiotic ranging from 0.008 to 8 *µ*g/mL. The first column of the 96-well plate contains the highest concentration of antibiotic and the 11th column contains the lowest concentration, with the 12th column containing no antibiotic, and 50 *µ*L of the diluted cell suspension was added to all wells. Plates were incubated at 37*^◦^*for 18 hours and examined for growth the next day. Results were only accepted if the observed MIC for the ATCC 25922 strain was within one doubling dilution of the expected result.

### Checkerboard minimum inhibitory concentration assay

Cultures of the *E. coli* MG1655 ancestral lineage and one of the six end-point isolates evolved in the presence of ibuprofen were prepared using a single colony inoculated into LB broth before incubation overnight at 37*^◦^*with agitation. Working stocks of ethidium bromide, ampicillin, ciprofloxacin, chloramphenicol, trimethoprim, colistin, and ibuprofen were prepared at four times the highest final concentration required by diluting in ISB. Overnight cultures were diluted 1:2000 in ISB. A 50 *µ*L aliquot of ISB was added to all columns of a 96-well plate, and 50 *µ*L working ibuprofen stock added to all wells of columns one and two. Starting with column two, the ibuprofen was serially diluted 1:2 across the plate up to and including column 11. A 50 *µ*L aliquot of one antibiotic working stock was then added to all wells of row A, the 1:2 dilution repeated down the plate up to and including row G, and 50 *µ*L removed from column 11 and row G before adding cells to keep the volume consistent. A 50 *µ*L sample of diluted overnight culture was then added to each well and mixed gently before the plate was covered and incubated for 18 hours at 37*^◦^*static. Plates were read following incubation and the presence or absence of growth noted.

### Statistical analyses

Area under the curve measurements were calculated using numpy.trapz in Python (v3.9.10). Significance testing was conducted using a one-way analysis of variance (ANOVA).

## Results

### Observed toxicity from pharmaceutical compounds against ***E. coli***

To first establish whether non-antibiotic pharmaceuticals can have observable toxicity, we screened a panel of compounds at a range of different concentrations (Table S1) against *E. coli* K-12 MG1655 as a model organism. Acetaminophen, ibuprofen, TiO_2_, propranolol, and metformin were all found to have significant negative effects on *E. coli* growth over a 24 hour incubation in comparison to the no-compound control at all concentrations tested (p < 0.05, one-way ANOVA, Fig. 1, Fig S1). The effect was predominantly noted as a reduction in the maximum OD reached. With the exception of TiO_2_ (where the highest concentration of 100 *µ*g/ml had a larger effect than other concentrations), altering the concentration of the compounds had little effect on the resulting growth kinetics. The non-antibiotic pharmaceuticals tested can therefore negatively impact growth of *E. coli* MG1655.

**Fig. 1.**
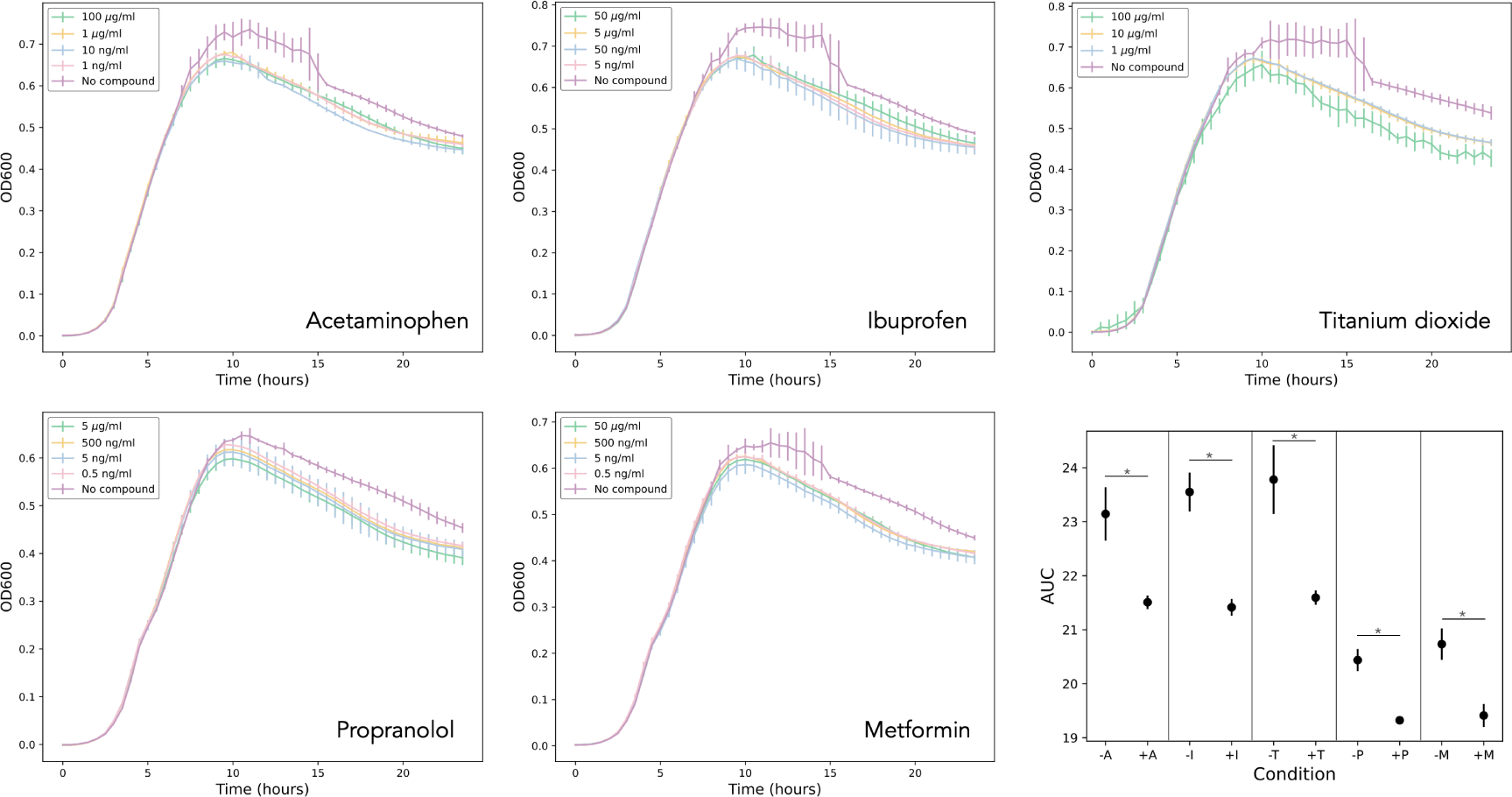
Toxicity screen of acetaminophen (+A), ibuprofen (+I), TiO_2_ (+T), metformin (+M), and propranolol (+P) at various concentrations against *E. coli* over a 24 incubation in a 96-well plate in a microplate reader. A no-compound control (‘No compound’, purple) was included for all screens. AUC values for the following concentrations are given against their representative no-compound control (-); 1 ng/mL acetaminophen, 5 ng/mL ibuprofen, 1 *µ*g/mL TiO_2_, 0.5 ng/mL propranolol, 0.5 ng/mL metformin. * p < 0.05, one-way ANOVA. All AUC values are shown in Fig. S1. Measurements in triplicate, error bars depict standard deviation.

### Upregulation in stress response and multidrug efflux genes in response to ibuprofen exposure

We noted a reduction in maximum OD following exposure to the compounds. To investigate the cause of this further, we conducted transcriptomic analysis on *E. coli* populations grown in the presence and absence of 50 *µ*g/mL ibuprofen. Ibuprofen was selected as exposure to this compound resulted in one of the larger reductions in maximum OD over a 24 hour time course, and it has been linked previously to a resistance phenotype by enhancing the transfer of resistance genes [31]. We found 16 genes were significantly upregulated in the presence of ibuprofen relative to the untreated control (Fig. 2). Those with the largest log fold change that could be influencing phenotype include *insC* (4.247), *nikA* (2.539), *yhcN* (1.396), *yhiM* (1.340), *lit* (1.289), and *mdtE* (1.285) (Table 1). NikA is a periplasmic binding protein for a nickel ATP-binding cassette (ABC) transporter. The *mdtE* gene encodes the membrane fusion protein component of a multidrug efflux system. Genes involved in a second multidrug efflux transporter, *emrA* and *emrD*, are also significantly upregulated, albeit to a lesser extent. The *yhcN* gene is linked to a response to stress, and *yhiM* to acid resistance. These data suggest that the observed reduction in maximum OD following ibuprofen exposure could be attributed to the cells undergoing a stress response or actively exporting the compound.

**Fig. 2.**
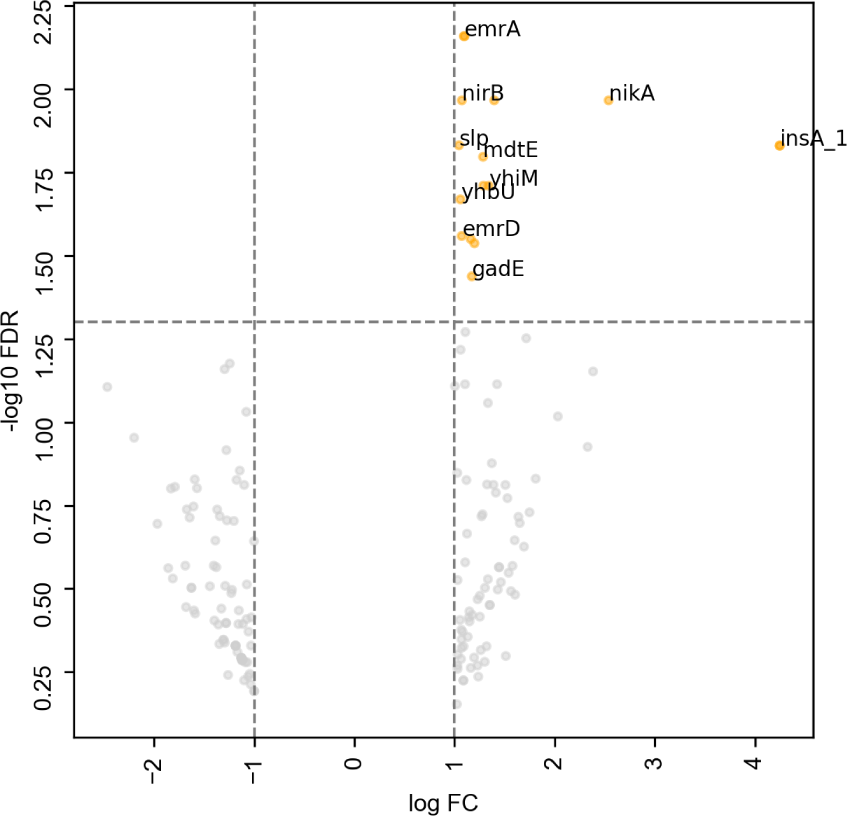
Genes significantly upregulated (yellow) (false discovery rate [FDR] threshold of p < 0.05 and an absolute log fold change [FC] of at least one) in the presence of ibuprofen. Genes which had an absolute log FC of at least one but did not reach the FDR threshold are shown in grey and are considered to not be significantly differentially expressed. Selected genes are labelled.

**Table 1.** *E. coli* genes upregulated significantly in the presence of ibuprofen, their function as assigned by Prokka, and the average (of biological triplicate) log fold change (FC) when normalised against *E. coli* grown in the absence of ibuprofen.

| Gene | Function | log FC |
| --- | --- | --- |
| *insA* | IS2 element protein | 4.247 |
| *insA* | IS2 element protein | 4.247 |
| *nikA* | Nickel ABC transporter - periplasmic binding protein | 2.539 |
| *yhcN* | Stress-induced protein | 1.396 |
| *yhiM* | Inner membrane protein with a role in acid resistance | 1.340 |
| *lit* | Cell death peptidase; phage exclusion; e14 prophage | 1.289 |
| *mdtE* | MdtEF-TolC multidrug efflux transport system - membrane fusion protein | 1.285 |
| *yhiD* | Putative Mg(2+) transport ATPase | 1.199 |
| *gadE* | DNA-binding transcriptional activator | 1.172 |
| *bhsA* | Outer membrane protein involved in copper permeability, stress resistance and biofilm formation | 1.161 |
| *aegA* | Putative oxidoreductase, Fe-S subunit | 1.099 |
| *emrA* | EmrAB-TolC multidrug efflux transport system - membrane fusion protein | 1.093 |
| *nirB* | Nitrite reductase, large subunit | 1.073 |
| *emrD* | Multidrug efflux transporter | 1.072 |
| *yhbU* | Putative peptidase (collagenase-like) | 1.059 |
| *slp* | Starvation lipoprotein | 1.044 |

### Co-exposure to ibuprofen and antibiotics does not alter MICs for *E. coli* MG1655

Microbial communities residing in or near industrial wastewater will be exposed to a cocktail of antibiotic and non-antibiotic compounds. The presence of a non-antibiotic pharmaceutical may induce a response that could alter the MIC of an antibiotic during co-exposure. To assess this, *E. coli* MG1655 and ATCC 25922 (as a control strain) were co-exposed to ibuprofen plus one of the following antimicrobial agents; ethidium bromide, ampicillin, ciprofloxacin, chloramphenicol, trimethoprim, and colistin. No change in MIC was observed for *E. coli* MG1655 for any ibuprofen-antimicrobial pair, with FIC scores indicating no synergy or antagonism (Table 2). This suggests that in laboratory strains of *E. coli*, co-exposure to ibuprofen alongside an antibiotic does not alter the resistance profile.

**Table 2.** MICs (mg/L) for six antimicrobial agents against the MG1655 ancestral lineage, the MG1655 strain evolved in the presence of ibuprofen, and an ATCC 25922 control strain (n=3, one biological replicate each, modal MIC value given with the exception of ciprofloxacin against MG1655 evolved where the median value is given). Checkerboard (CB) MICs (mg/L) for *E. coli* MG1655 and ATCC 25922 co-exposed to ibuprofen plus an antimicrobial agent, one of; ethidium bromide, ampicillin, ciprofloxacin, chloramphenicol, trimethoprim, or colistin (n=3, one biological replicate each, modal MIC value given). Fractional Inhibitory Concentration (FIC) scores calculated whereby 0.5 – 4 indicates no synergy or antagonism. - indicates not applicable/not tested.

| Agent | MIC ancestor | MIC evolved | MIC ATCC 25922 | CB MIC ancestor | FIC ancestor | CB MIC ATCC 25922 | FIC ATCC 25922 |
| --- | --- | --- | --- | --- | --- | --- | --- |
| Ethidium bromide | 5 | 512 | 256 | 512 | 2 | 256 | 2 |
| Ampicillin | 4 | 8 | 8 | 8 | 3 | 8 | 2 |
| Ciprofloxacin | 0.0156 | 0.0156 | 0.0156 | 0.0156 | 2 | 0.0156 | 2 |
| Chloramphenicol | 8 | 8 | 4 | 8 | 2 | 4 | 2 |
| Trimethoprim | 0.25 | 0.25 | 0.25 | 0.5 | 3 | 0.5 | 3 |
| Colistin | 4 | 4 | 4 | 2 | 1.5 | 4 | 2 |
| Ibuprofen | >200 | - | >200 | >200 | - | - | - |

### Long-term exposure to non-antibiotic pharmaceuticals impacts *E. coli* growth but does not select for **cross-resistance to antibiotics**

After establishing the negative impact on *E. coli* growth in the presence of selected non-antibiotic pharmaceuticals, we investigated whether this would be sufficient to act as a selective pressure during long-term exposure. We therefore propagated populations in the presence and absence of acetaminophen, ibuprofen, TiO_2_, metformin, and propranolol individually, passaging cells every 24 hours for 30 days. The evolved populations were then screened in the presence and absence of their selection compound to assess growth in comparison to the ancestral lineage. We observed a decrease in the maximum OD readings reached by the evolved populations in their selection media in comparison to the ancestral lineage, suggesting that prolonged exposure to the compounds did not select for improved growth (Fig. 3). This difference was statistically significant in the majority of populations (p<0.05, one-way ANOVA, Fig. S2), and was most notable for the populations exposed to ibuprofen and TiO_2_ (Fig. 3). The reduction in OD observed in the no compound control was calculated to be not significant (Fig. S2). The difference remained when the populations were exposed to 100x concentration of the compounds (Fig. S3), and when all populations were passaged through a seven-day ‘recovery’ experiment in NB with no added pharmaceutical (Fig. S4). This suggests therefore that the growth patterns observed in the evolved populations was not a transient, reversible effect as a result of long-term toxicity.

**Fig. 3.**
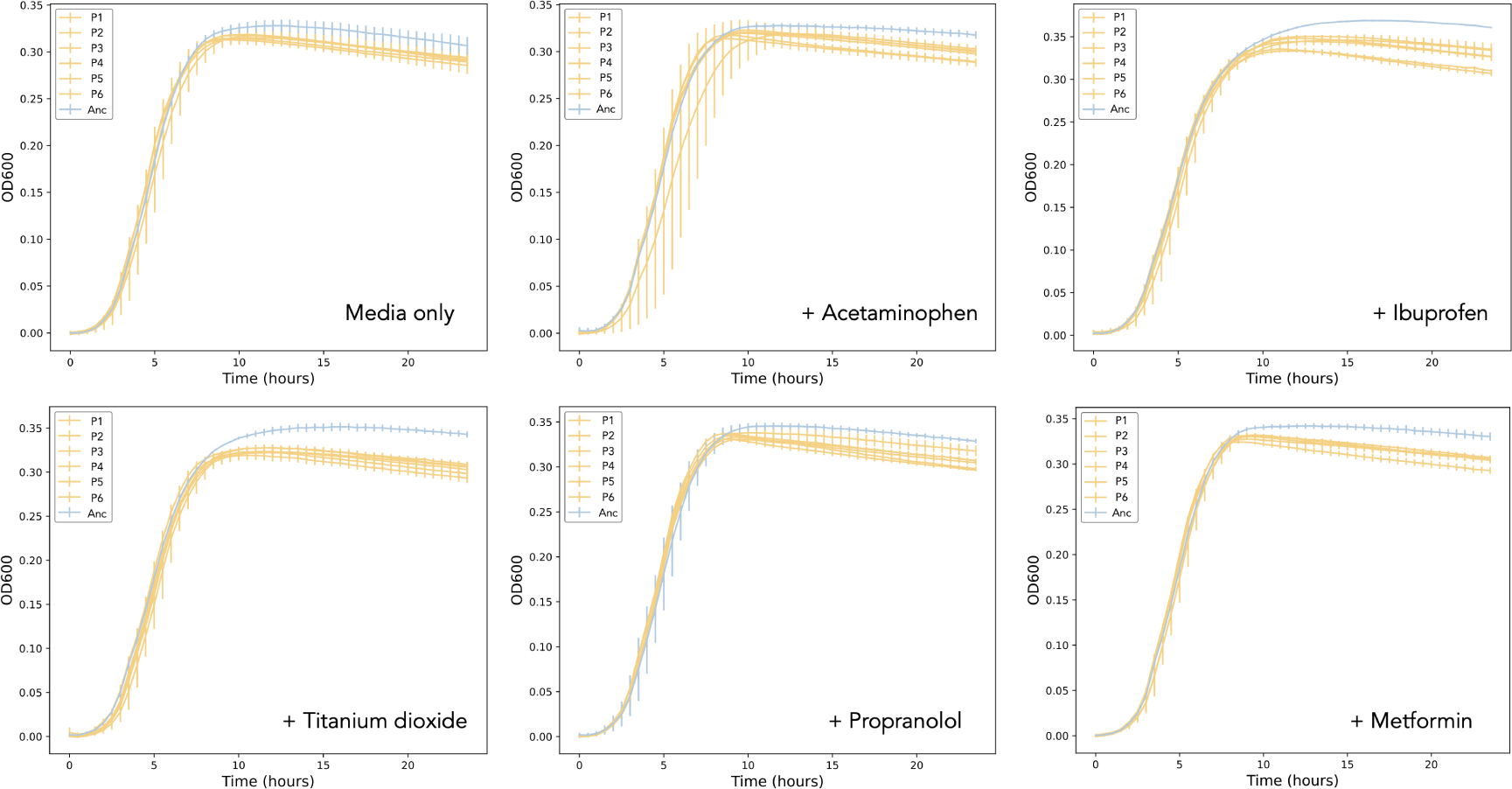
Growth of evolved populations (yellow, six independent biological replicates P1-P6) in the presence of the compound in which their selection experiment was conducted in comparison to growth of the ancestral lineage (blue, Anc); (clockwise from top left) media-only control, acetaminophen, ibuprofen, metformin, propranolol, and TiO_2_. Measurements in triplicate, error bars depict standard deviation.

To establish whether there were any significant single nucleotide polymorphisms (SNPs) arising as a result of the selection experiments, short-read assemblies were generated for three replicates from each of the control and test conditions and the sequences analysed using Snippy. No mutations parallel between independent evolving populations were found within treatments (Table S3). SNPs in *recQ* (ATP-dependent DNA helicase) and *ygeA* (a putative racemase) were observed in single replicates of evolved populations exposed to acetaminophen. The singular occurrence of each suggests they arose as a result of drift rather than selection, or that selection was not strong enough for them to reach fixation within the other populations. An analysis of the gene presence/absence patterns across the hybrid assembled evolved isolate genomes highlighted sequence variation in *ldrA* gene in three sequences; one control, and one each evolved in acetaminophen and TiO_2_. All had three SNPs compared to the ancestral MG1655 (Table S2). The TiO_2_ and the control had identical SNPs, whereas the three SNPs in the acetaminophen-exposed isolate were different. Again, these variations occurred in a single replicate per condition only.

The IS*2* element *insA* was shown to be upregulated in the presence of ibuprofen. IS element transposition could be a cause of the differences in growth patterns between the ancestral and evolved isolates. To assess this, hybrid assemblies were generated and the distribution of IS elements then established using ISEScan and ISFinder. All IS elements were present in equal numbers between the ancestor and all evolved lineages. IS*2*, IS*30*, and IS*1* elements were interrogated in depth and showed no evidence of movement between any evolved isolate and the ancestor. Overall, these data suggest that although the presence of the non-antibiotic pharmaceuticals has a negative effect on the growth on *E. coli*, they do not exert a selection pressure during prolonged exposure at the timescale and concentrations tested.

Given the observed upregulation in known efflux systems following ibuprofen treatment, we examined whether prolonged exposure to ibuprofen may select for cross-resistance to antibiotics. The MICs of several antimicrobials were analysed for a strain evolved in the presence of ibuprofen compared to the ancestral isolate. Ethidium bromide, ampicillin, ciprofloxacin, chloramphenicol, trimethoprim, and colistin were chosen based on the previously mentioned transcriptomic data as compounds which might logically have altered MICs as a result of ibuprofen exposure. The MICs of the evolved isolate for all antibiotics tested were either the same or within one doubling dilution of the ancestor (Table 2). This indicates the long-term exposure to ibuprofen, regardless of the differential gene expression, does not co-select for resistance to the antibiotics tested.

## Discussion

The evolution of bacteria, including *E. coli*, in response to exposure to antibiotics is well understood. The effects of short- and long-term exposure to non-antibiotic pharmaceuticals is however less clear. This includes their potential toxicity and the likelihood that their presence might select for cross-resistance to antibiotics. With increasing evidence for the presence of non-antibiotic pharmaceuticals in proximity to production plants [5; 18], it is becoming apparent that these are compounds that require further investigation into their potential to impact local microbial populations. To begin to address this, we screened a panel of five non-antibiotic pharmaceuticals against a laboratory strain of *E. coli* to assess their potential toxicity and found that all five, to various degrees, reduced the maximum OD reached by the population over a 24 hour incubation. This therefore suggests that these compounds may, even at low concentrations, have a negative effect on members of the microbial communities surrounding production plants. Our results contribute to published work on ibuprofen toxicity, whereby it was demonstrated through disc diffusion assays to not have antimicrobial activity against *E. coli* [30].

We then uncovered 16 genes that are upregulated significantly when *E. coli* was grown in the presence of ibuprofen. Notable amongst these genes were two periplasmic adaptor proteins from different multidrug efflux systems; *mdtE* and *emrA*. MdtEF-TolC is a resistance nodulation division (RND) family pump with beta-lactams, benzalkonium chloride, macrolides, and oxazolidinones as known substrates, and EmrAB-TolC is a major facilitator superfamily (MFS) efflux pump that confers resistance to compounds including fluoroquinolones in *E. coli* [50]. Efflux pump expression is often upregulated in the presence of toxic compounds to prevent their accumulation inside the cell [51; 52; 53; 54]. There is some existing evidence linking efflux pumps to a response to non-antibiotic pharmaceuticals. Exposing *S. aureus* to diclofenac has been shown to downregulate a putative *emrAB*-family pump [55], which contrasts the upregulation we observed in *E. coli* following ibuprofen exposure. The response could therefore be specific to the species, compound, or pump, and more work is needed to unravel this potential interaction.

Ibuprofen is an example of a partial proton motive force (PMF) uncoupler that can inhibit the function of RND and MFS pumps [56]. The nitrite reductase *nirB* was identified as another significantly upregulated gene. Nitrate reduction has been shown to enhance bacterial survival in the presence of agents that dissipate PMF [55]. This therefore provides tentative support to a hypothesis that *E. coli* may be using nitrate reduction to ameliorate the dissipation of PMF in the presence of ibuprofen, enabling the function of the RND and MFS pumps.

Ibuprofen exposure also resulted in the upregulation of several genes linked to a response to stress, including *yhcN*, the inner membrane protein *yhiM*, and the outer membrane protein *bhsA*. Existing research has shown a *bhsA* mutant of *E. coli* to be more sensitive to a variety of stressors including acid [57], and *yhcN* has been linked to cytoplasm pH stress [58]. Additionally, the gene observed to be upregulated to the greatest extent in our data was *insA*, and IS*2* and other IS elements are known to be upregulated in response to stress [59; 60]. An upregulation of stress response genes in response to non-antibiotic pharmaceuticals has been noted previously in *A. baylyi*, where the uptake of antibiotic resistance genes was shown to be facilitated by the presence of compounds including ibuprofen and propranolol [31]. There, analysis including transcriptomics linked the observation to increased stress and the over-production of reactive oxygen species, amongst other characteristics.

Our data suggest that the non-antibiotic components within pharmaceutical production waste may affect the local microbial communities, as over time the toxicity observed here may deplete species or genera within the communities, altering their composition. Despite the observed reduction in maximum OD, we found that prolonged exposure to this panel of non-antibiotic pharmaceuticals did not result in significant genetic changes across multiple independent populations. It is possible that the stress induced by the compounds was dealt with sufficiently by, for example, the upregulation of efflux pumps, thereby reducing the selection pressure, or that the reduced OD was not due to changes in carrying capacity but rather due to morphological changes following induction of stress responses. We also found no evidence of synergy when the ancestral strain was co-exposed to ibuprofen and one of a panel of antibiotics. Additionally, when the same panel of antibiotics were tested against the ancestor and the strain evolved in the presence of ibuprofen, there was no evidence of altered MICs in the latter that would indicate selection for cross-resistance to antibiotics. This is a reassuring initial investigation given the large quantities of pharmaceutical production waste entering local ecosystems. Whilst the panel of non-antibiotic pharmaceuticals tested here is small, they are commonly used and consistently found in water bodies, and therefore can be considered a representative sample [61; 62; 63; 64]. Acetaminophen, ibuprofen, metformin, and propranolol have also been identified as priority pharmaceuticals in India [65]. The use of a standard laboratory strain of *E. coli* is a useful starting point, but previous work suggesting that the activity of non-antibiotic pharmaceuticals may be strain-specific [21] underscores the need for broad spectrum testing before definitive conclusions can be drawn. Variations in concentrations over time as a result of effluent changes, dry seasons, and climate change should also be considered, and there is a need to extend this research to encompass production waste as a holistic entity against environmentally relevant populations.

## Acknowledgements

Illumina genome sequencing was provided by MicrobesNG (www.microbesng.com). Standard RNA sequencing was performed by GENEWIZ from Azenta Life Sciences. RJH was supported by a NERC grant (NE/T01301X/1) awarded to AM, LJC and IA.

## Data availability statement

The datasets generated and analysed during the current study are available from NCBI BioProject with accession PRJNA1005239.

## Author contributions

RJH: methodology, formal analysis, investigation, data curation, writing - original draft, visualization. AES: investigation, data curation. SJE: methodology, investigation, vizualisation. RAM: methodology. HS: validation, investigation. EAC: methodology. MJB: methodology. JMAB: conceptualization, resources, supervision. IA: funding acquisition. LJC: conceptualization, funding acquisition. AM: conceptualization, methodology, supervision, funding acquisition. All authors: writing - review & editing.

## Supplementary

**Fig. S1.**
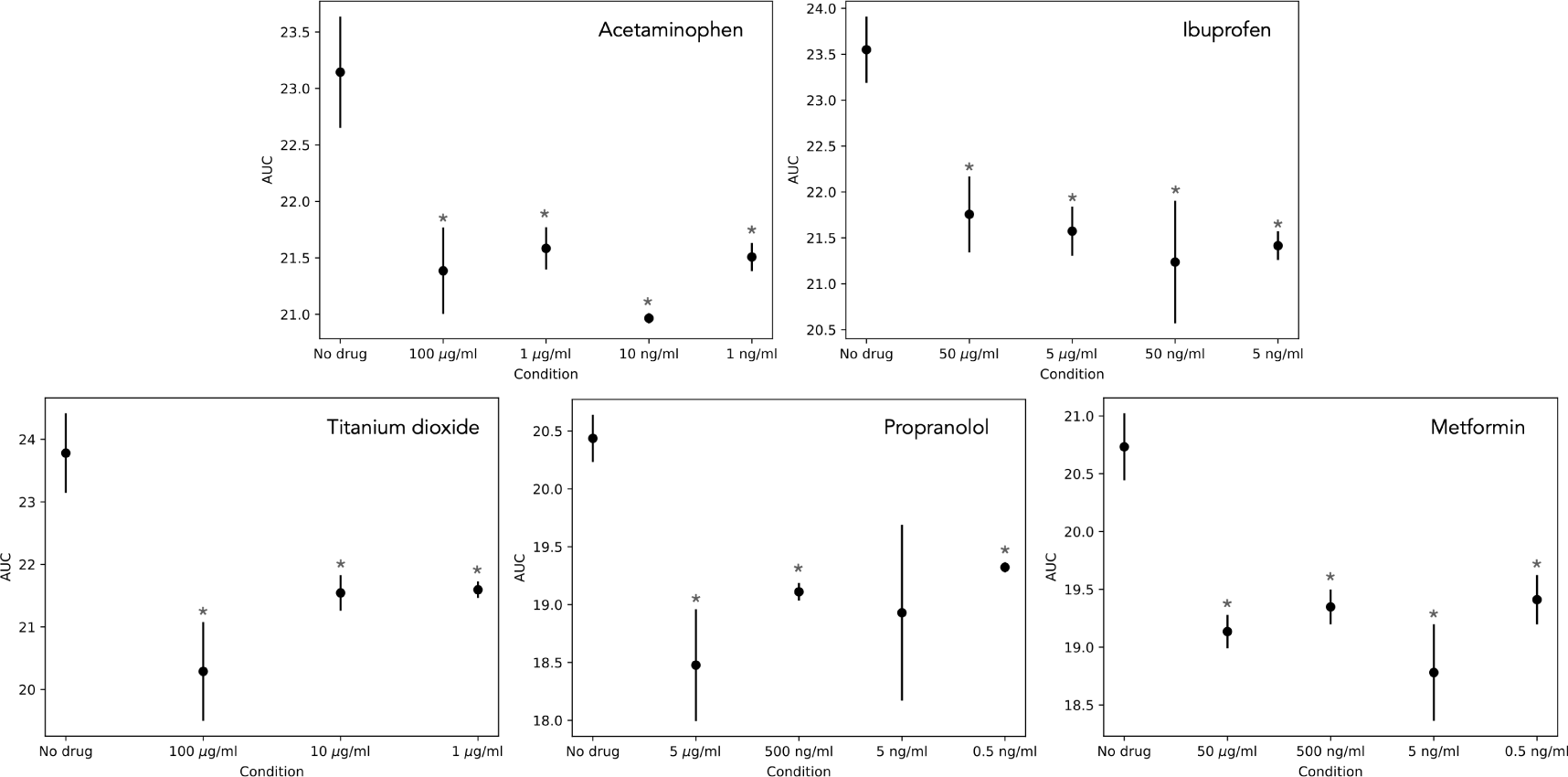
AUC values for a toxicity screen of (clockwise from top left) acetaminophen, ibuprofen, metformin, propranolol, and TiO_2_ at various concentrations against *E. coli*, relating to Fig. 1. A no-compound control (‘No drug’) was included for each screen. * p < 0.05, one-way ANOVA. Measurements in triplicate, error bars depict standard deviation.

**Fig. S2.**
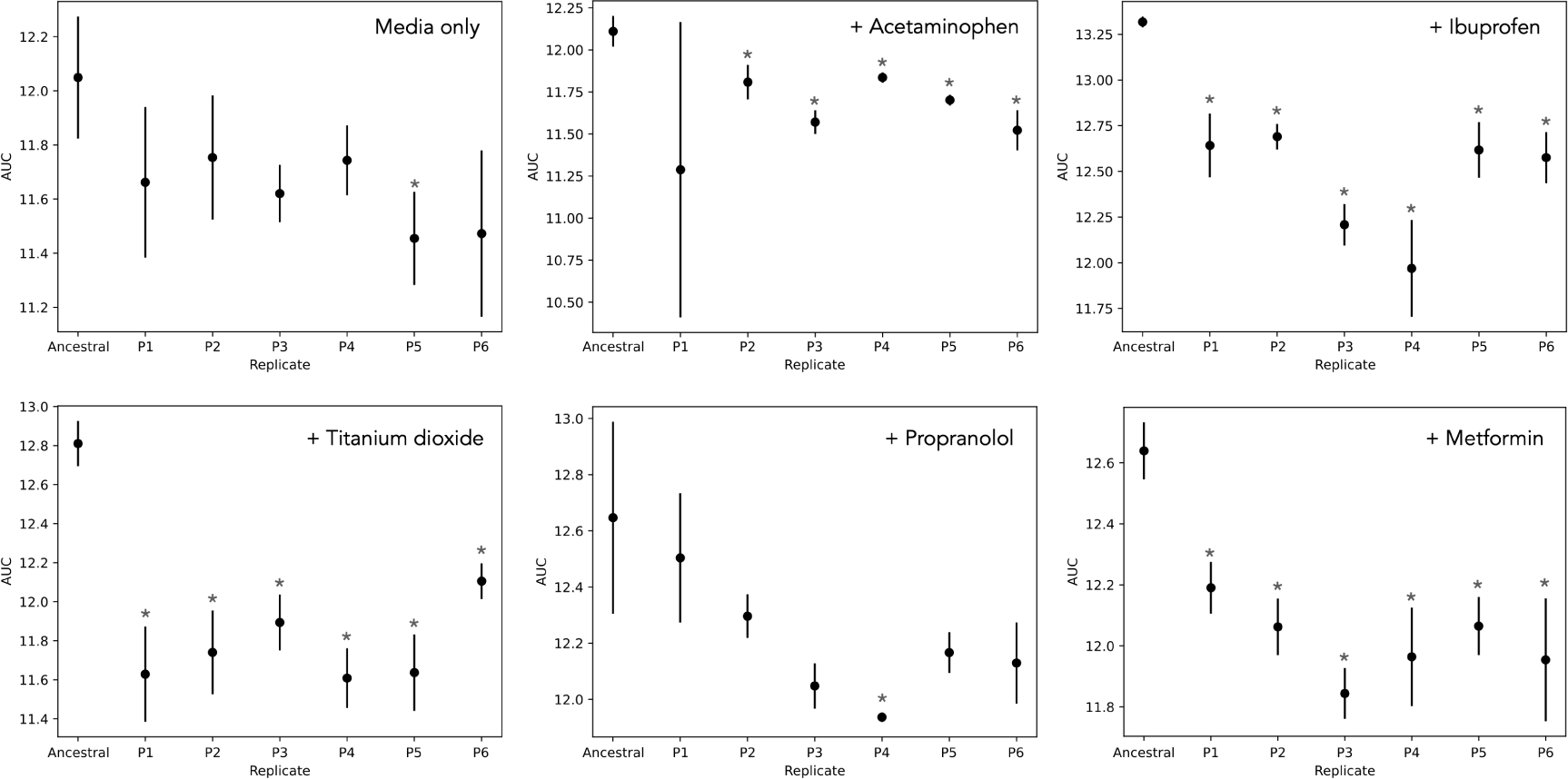
AUC values for evolved populations (six independent biological replicates P1-P6) in a fresh sample of their respective evolution media in comparison to the ancestral lineage, relating to Fig. 3. * p < 0.05, one-way ANOVA. Measurements in triplicate, error bars depict standard deviation.

**Fig. S3.**
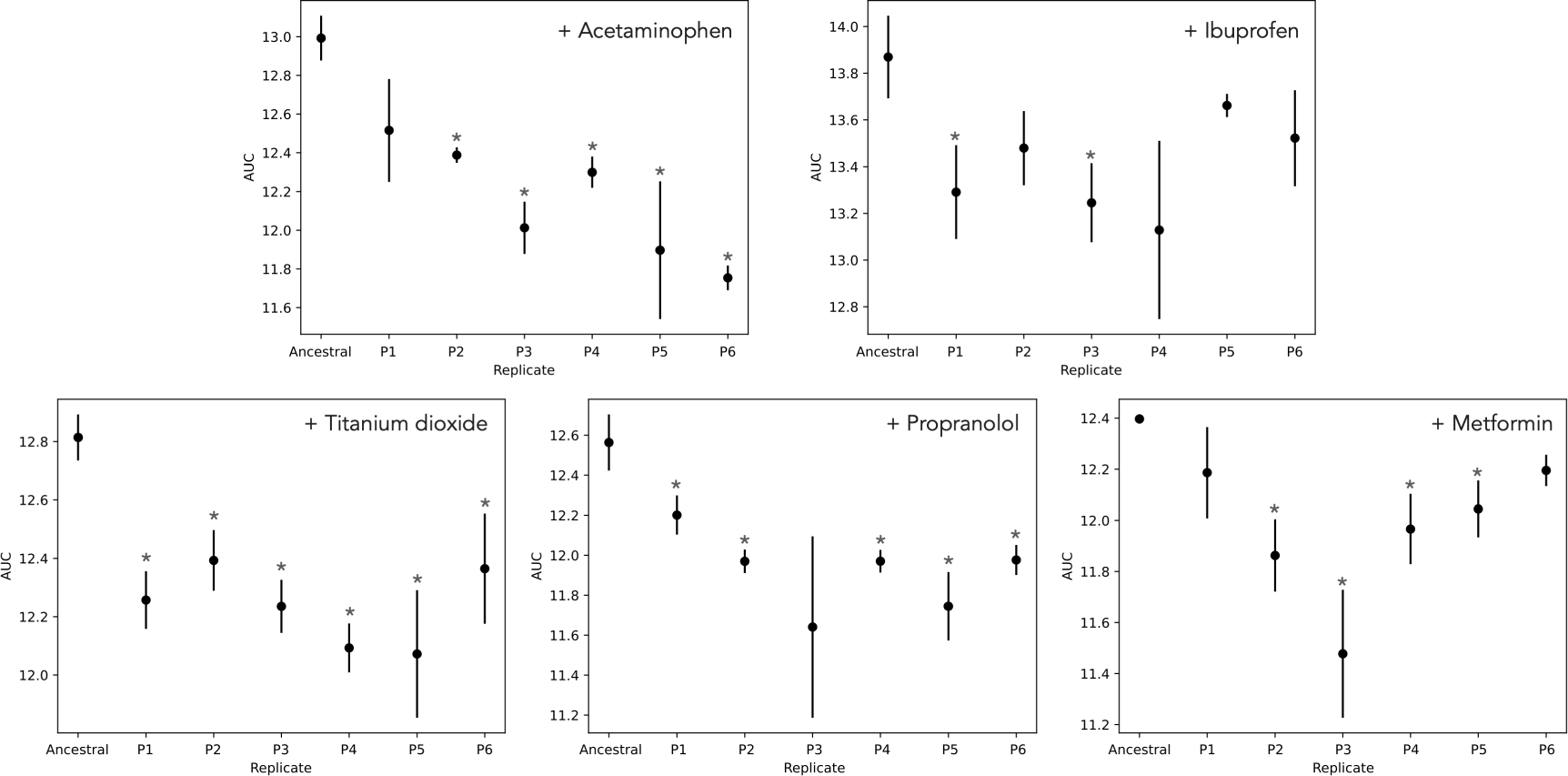
AUC values for evolved populations (six independent biological replicates P1-P6) in the presence of 100x concentration of the compound in which their selection experiment was conducted in comparison to growth of the ancestral lineage. * p < 0.05, one-way ANOVA. Measurements in triplicate, error bars depict standard deviation.

**Fig. S4.**
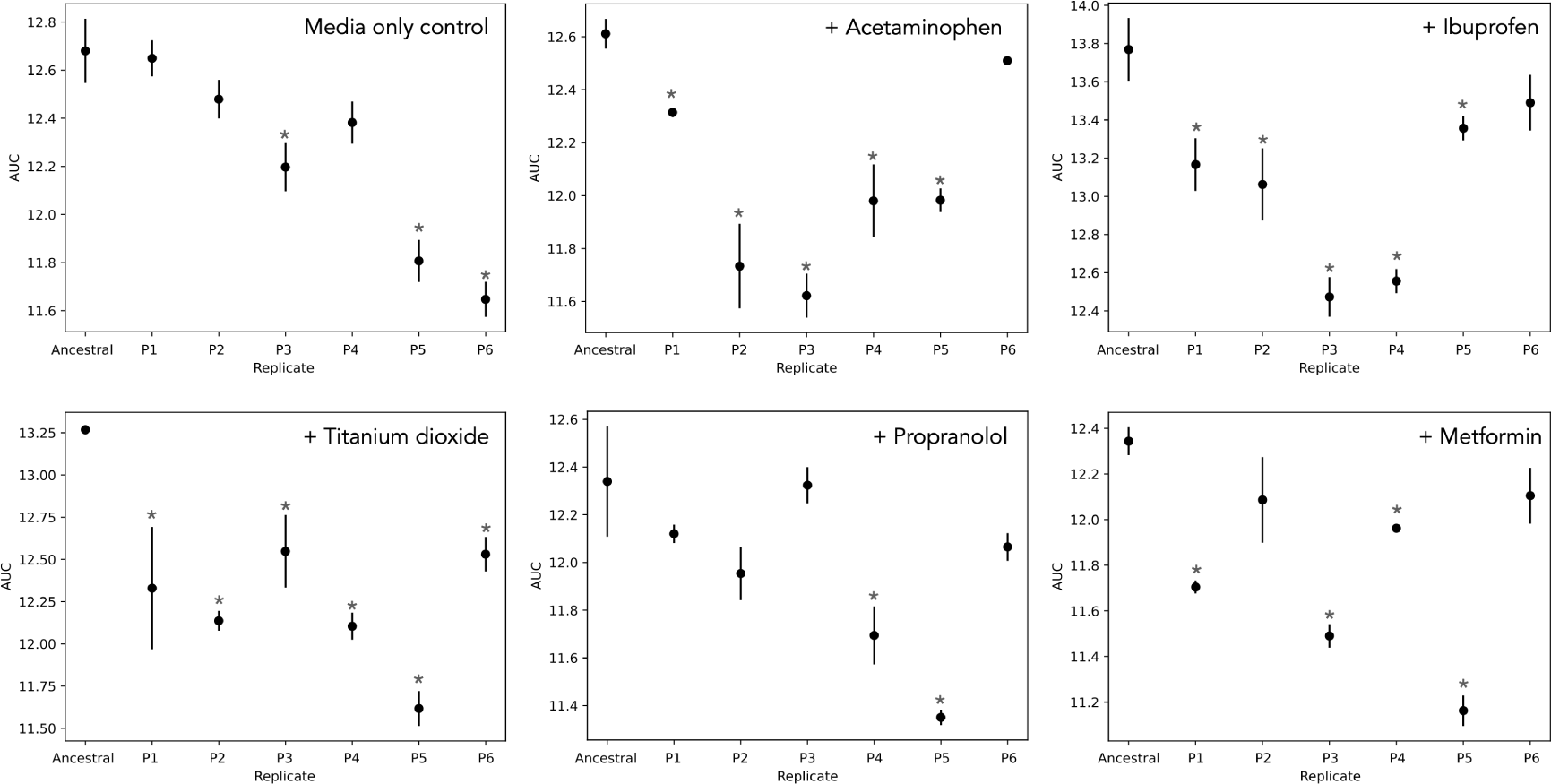
AUC values for evolved populations (six independent biological replicates P1-P6) following serial passaging for seven days in the absence of pharmaceuticals in comparison to the ancestral lineage. * p < 0.05, one-way ANOVA. Measurements in triplicate, error bars depict standard deviation.

**Table S1.**
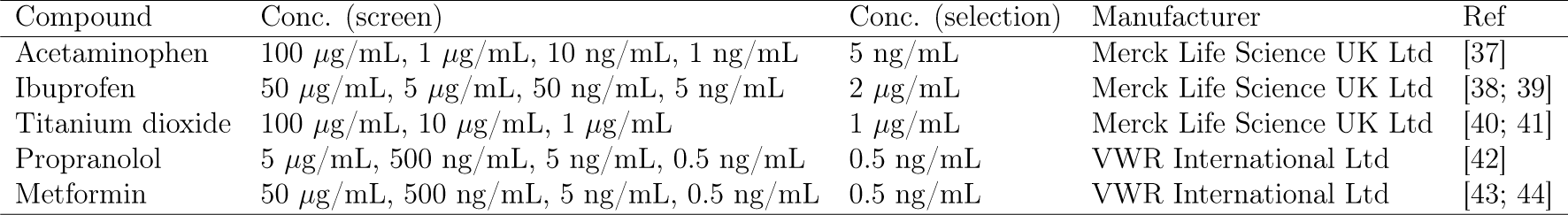
Concentrations used to screen the non-antibiotic pharmaceutical compounds for toxicity against *E. coli*, and the final concentration chosen for selection experiments. No-compound controls were also used as comparisons for all screens. Relevant references are also included.

| Compound | Conc. (screen) | Conc. (selection) | Manufacturer | Ref |
| --- | --- | --- | --- | --- |
| Acetaminophen | 100 µg/mL, 1 µg/mL, 10 ng/mL, 1 ng/mL | 5 ng/mL | Merck Life Science UK Ltd | [37] |
| Ibuprofen | 50 µg/mL, 5 µg/mL, 50 ng/mL, 5 ng/mL | 2 µg/mL | Merck Life Science UK Ltd | [38; 39] |
| Titanium dioxide | 100 µg/mL, 10 µg/mL, 1 µg/mL | 1 µg/mL | Merck Life Science UK Ltd | [40; 41] |
| Propranolol | 5 µg/mL, 500 ng/mL, 5 ng/mL, 0.5 ng/mL | 0.5 ng/mL | VWR International Ltd | [42] |
| Metformin | 50 µg/mL, 500 ng/mL, 5 ng/mL, 0.5 ng/mL | 0.5 ng/mL | VWR International Ltd | [43; 44] |

**Table S2.** SNP differences observed in the *ldrA* gene in the hybrid assemblies of the evolved isolates in comparison to the ancestor. R = test replicate, SNP = single nucleotide polymorphism.

| Condition | SNP | Position |
| --- | --- | --- |
| Control R3 | T→C | 28 |
|  | A→G | 47 |
|  | A→G | 87 |
| + Acetaminophen R3 | T→C | 24 |
|  | T→C | 64 |
|  | A→G | 83 |
| + Titanium dioxide R3 | T→C | 28 |
|  | A→G | 47 |
|  | A→G | 87 |

**Table S3.** SNPs identified in each sequence (R = test replicate) as predicted by Snippy. The nucleotide(s) in the reference (ref) and comparison (alt) sequences, the locus tag of the gene, and its product are given.

| Condition | Ref | Alt | Locus tag | Gene | Product |
| --- | --- | --- | --- | --- | --- |
| **Ancestor** | A | C | 00551 | *rhcC* | Protein RhsC |
| **Media only R1** | A | G | 00431 |  | Uncharacterized protein HI_1672 |
|  | C | G | 00553 |  | Hypothetical protein |
|  | A | G | 00553 |  | Hypothetical protein |
|  | T | G |  |  |  |
| **Media only R2** | A | C | 00551 | *rhcC* | Protein RhsC |
| **Media only R3** | A | C | 00551 | *rhcC* | Protein RhsC |
|  | A | C | 03981 | *sspA* | Stringent starvation protein A |
| **+Acetaminophen R1** | A | C | 00551 | *rhcC* | Protein RhsC |
|  | C | G | 00553 |  | Hypothetical protein |
|  | A | G | 00553 |  | Hypothetical protein |
|  | C | A |  |  |  |
|  | T | G | 03610 | *recQ* | ATP-dependent DNA helicase |
| **+Acetaminophen R2** | A | C | 00551 | *rhcC* | Protein RhsC |
|  | C | G | 00553 |  | Hypothetical protein |
|  | A | G | 00553 |  | Hypothetical protein |
| **+Acetaminophen R3** | C | G | 00553 |  | Hypothetical protein |
|  | A | G | 00553 |  | Hypothetical protein |
|  | GCGCC | G | 01387 | *ygeA* | Putative racemase |
| **+Ibuprofen R1** | A | C | 00551 | *rhcC* | Protein RhsC |
| **+Ibuprofen R2** | C | G | 00553 |  | Hypothetical protein |
|  | A | G | 00553 |  | Hypothetical protein |
| **+Ibuprofen R3** | A | C | 00551 | *rhcC* | Protein RhsC |
|  | A | G | 00553 |  | Hypothetical protein |
| **+Titanium dioxide R1** | A | C | 00551 | *rhcC* | Protein RhsC |
|  | C | G | 00553 |  | Hypothetical protein |
|  | A | G | 00553 |  | Hypothetical protein |
| **+Titanium dioxide R2** | A | C | 00551 | *rhcC* | Protein RhsC |
|  | A | G | 00553 |  | Hypothetical protein |
| **+Titanium dioxide R3** | A | C | 00551 | *rhcC* | Protein RhsC |
|  | C | G | 00553 |  | Hypothetical protein |
|  | A | G | 00553 |  | Hypothetical protein |
| **+Propranolol R1** | A | C | 00551 | *rhcC* | Protein RhsC |
|  | T | G |  |  |  |
| **+Propranolol R2** | A | C | 00551 | *rhcC* | Protein RhsC |
| **+Propranolol R3** | A | C | 00551 | *rhcC* | Protein RhsC |
|  | A | G | 00553 |  | Hypothetical protein |
|  | G | A |  |  |  |
| **+Metformin R1** | A | C | 00551 | *rhcC* | Protein RhsC |
|  | C | G | 00553 |  | Hypothetical protein |
|  | A | G | 00553 |  | Hypothetical protein |
| **+Metformin R2** | A | C | 00551 | *rhcC* | Protein RhsC |
|  | A | G | 00553 |  | Hypothetical protein |
|  | G | T |  |  |  |
| **+Metformin R3** | None |  |  |  |  |

